# Clinical evaluation of the Multimapping technique for simultaneous myocardial T_1_ and T_2_ mapping

**DOI:** 10.1101/2022.06.02.494576

**Authors:** Charlotta Jarkman, Carl-Johan Carlhäll, Markus Henningsson

**Author notes:** **Correspondence:** Markus Henningsson. Equal contributors (Carl-Johan Carlhäll, Markus Henningsson).

## Abstract

The Multimapping technique was recently proposed for simultaneous myocardial T_1_ and T_2_ mapping. In this study we evaluate its correlation with clinical reference mapping techniques in patients with a range of cardiovascular diseases (CVD), compare image quality and inter- and intra-observer repeatability. Multimapping consists of a ECG-triggered, 2D single-shot bSSFP readout with inversion recovery and T_2_ preparation modules, acquired across 10 cardiac cycles. The sequence was implemented at 1.5T and compared to clinical reference mapping techniques, Modified Look-Locker inversion recovery (MOLLI) and T_2_ prepared bSSFP with four echo times (T_2_bSSFP), and compared in 47 patients with CVD (of which 44 were analyzed). In diseased myocardial segments (defined as presence of late gadolinium enhancement) there was a high correlation between Multimapping and MOLLI for native myocardium T_1_ (r^2^=0.73), ECV (r^2^=0.91) and blood T_1_ (r^2^=0.88), and Multimapping and T_2_bSSFP for native myocardial T_2_ (r^2^=0.80). In healthy myocardial segments a bias for native T_1_ (Multimapping=1116±21 ms, MOLLI=1002±21, *P*<0.001), post-contrast T_1_ (Multimapping=479±31 ms, MOLLI=426±27 ms, 0.001), ECV (Multimapping=21.5±1.9%, MOLLI=23.7±2.3%, *P*=0.001) and native T_2_ (Multimapping=48.0±3.0 ms, T_2_bSSFP=53.9±3.5 ms, *P*<0.001) was observed. The image quality for Multimapping was scored as higher for all mapping techniques (native T_1_, post-contrast T_1_, ECV and T_2_bSSFP) compared to the clinical reference techniques. The inter- and intra-observer agreement was excellent (intraclass correlation coefficient, ICC>0.9) for most measurements, except for inter-observer repeatability of Multimapping native T_1_ (ICC=0.87), post-contrast T_1_ (ICC=0.73) and T_2_bSSFP native T_2_ (ICC=0.88). Multimapping show high correlations with clinical reference mapping techniques for T_1_, T_2_ and ECV in a diverse cohort of patients with different cardiovascular diseases. Multimapping enables simultaneous T_1_ and T_2_ mapping and can be performed in a short breath-hold, with image quality superior to that of the clinical reference techniques.

## 1 Introduction

Myocardial T_1_ and/or T_2_ values are altered in many cardiovascular diseases (1). T_1_ and T_2_ quantification, along with disease-specific patterns of regional and global distribution, can be captured with myocardial mapping techniques (2). In the last 15-20 years a number of T_1_ and T_2_ mapping techniques have been published, with different strengths and weaknesses in terms of quantification accuracy, precision, scan time, spatial resolution and coverage (3). Despite being one of the first T_1_ mapping techniques, the modified Look-Locker inversion recovery (MOLLI) remains the most clinically used method due to its high precision and availability on all major scanner platforms (4,5). However, MOLLI T_1_ accuracy is relatively low and the quantification is susceptible to confounding effects from heart-rate or T_2_-dependent variability, magnetization transfer effects, motion artifacts and system imperfections (5,6). Different T_1_ mapping methods have been proposed to address these shortcomings, yet have failed to make significant inroads in the market share of clinical use (7–11). Myocardial T_2_ mapping can be performed with multi-echo spin-echo or T_2_-prepared balanced steady-state free precession (T_2_bSSFP) techniques (12–14). The latter approach is likely the most widely used clinically due to its relative robustness to physiological motion.

In recent years, there has been a growing interest in techniques to simultaneously map T_1_ and T_2_ in a single scan (15–21). Advantages of this approach compared to conventional mapping, which is performed separately for T_1_ and T_2_, is that the images are intrinsically spatially aligned, scan time is typically shorter and the confounding effects of T_1_ or T_2_ on the quantification of the opposite parameter is minimized. Despite the many theoretical advantages of simultaneous T_1_ and T_2_ mapping, there is a paucity of translational studies using these techniques in patients with cardiovascular disease (22,23). This may be due to the more sophisticated acquisition, reconstruction and mapping strategies necessary for such techniques, which pose challenges for clinical translation. Recently, a new technique for simultaneous T_1_ and T_2_ mapping, termed Multimapping, was proposed using a standard Cartesian trajectory and evaluated (primarily) in healthy subjects (24). Due to its simplicity Multimapping may be readily applied in a clinical setting to enable evaluation of simultaneous T_1_ and T_2_ mapping in patients with cardiovascular disease.

The primary aim of this study is to validate Multimapping T_1_ and T_2_ values against clinical reference techniques in patients with different cardiovascular diseases in terms of parameter quantification and image quality. A secondary study aim is to evaluate Multimapping intra- and inter-observer variability.

## 2 Materials and Methods

### 2.1 Study population

All patients provided written informed consent prior to participation and the study was approved by the local ethics committee (Linköping Regional Ethics Committee, 2015/396–31) and conducted according to the Declaration of Helsinki. Patients referred for CMR at Linköping University Hospital between June and November 2021 were considered for inclusion in this study. In total, 47 patients were recruited. Datasets from three patients were excluded, two because no late gadolinium enhancement (LGE) images were acquired and one due to excessive fold-over artifacts. Clinical characteristics of the remaining patients can be seen in Table 1. Of the included patients, normal cardiac MRI scan was found in 15 (34.1%) patients, myocarditis in 11 (25%) patients, dilated cardiomyopathy (DCM) in 6 (13.6%) patients, ventricular hypertrophy (hypertrophic cardiomyopathy or hypertrophy of unknown origin) in 5 (11.4%) patients, ischemic myocardial injury (acute/recent or old) in 3 (6.8%) patients, arrhythmogenic right ventricular cardiomyopathy in 2 (4.5%) patients, pericarditis in 1 (2.3%) patient and congenital heart disease in 1 (2.3%) patient.

**Table 1.**
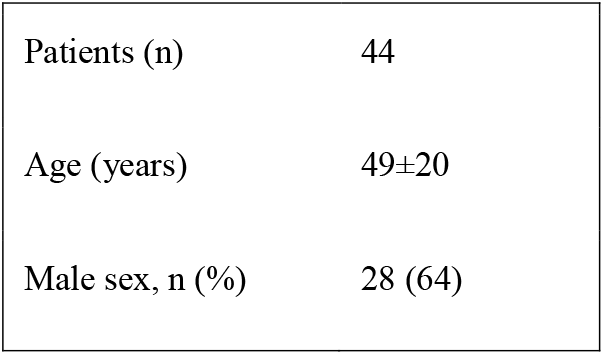
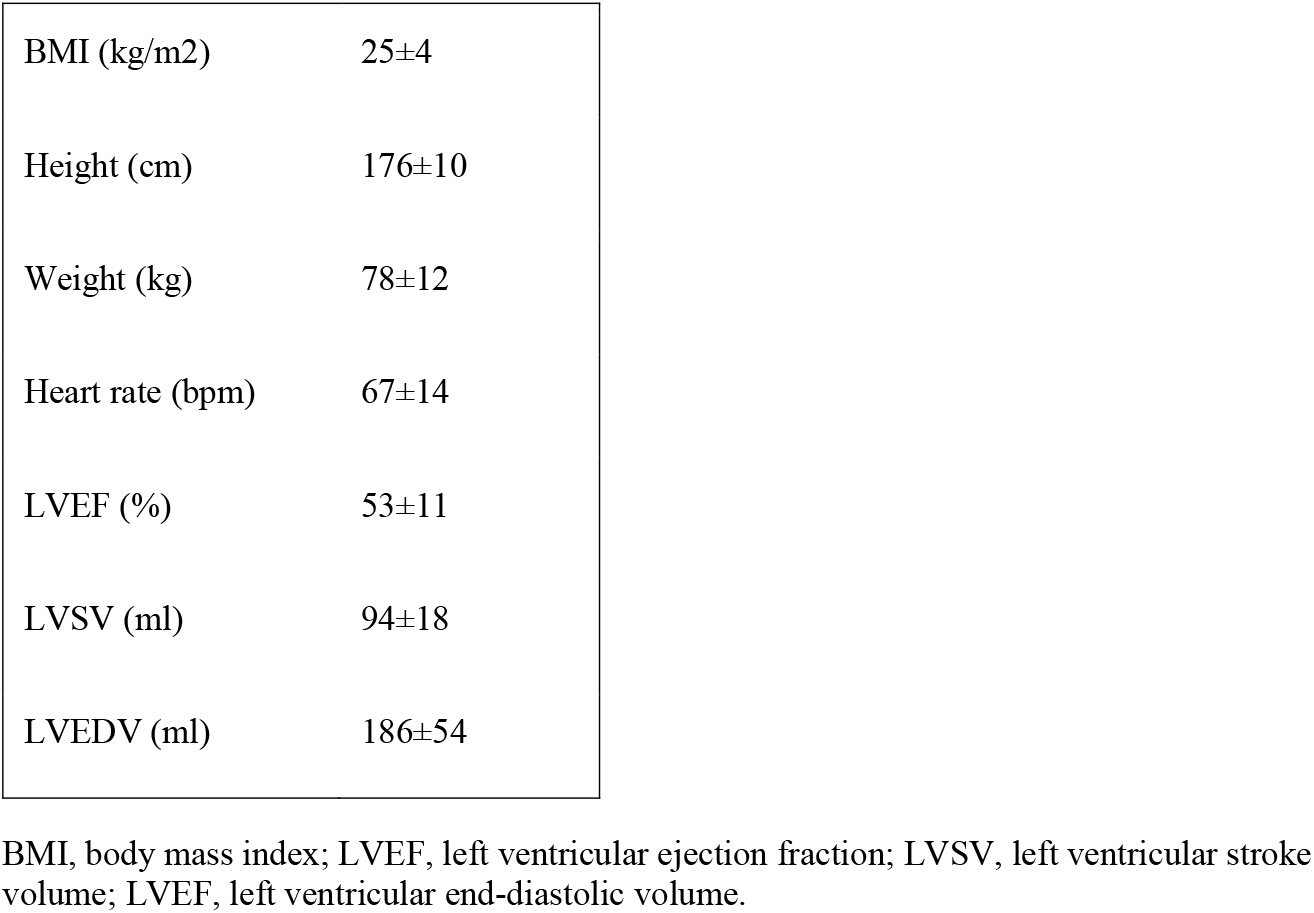
Clinical Characteristics

### 2.2 Data acquisition and reconstruction

All scans were performed on a 1.5T Philips clinical CMR scanner (Philips Healthcare, Best, The Netherlands) using a 28-channel torso coil. The Multimapping pulse sequence and post-processing steps are illustrated in Figure 1. Ten single-shot images are acquired across consecutive cardiac cycles using balanced-steady state free precession (bSSFP) readouts, triggered to the mid-diastolic rest period. Adiabatic inversion radiofrequency (RF) pulses with delay times of 300 ms are performed in the 1^st^ and 5^th^ cardiac cycle to improve T_1_ sensitization. The inversion pulse used a hyperbolic secant shape, had a duration of 8.4 ms and a B_1_ amplitude of 13.5 µT. A previous study has shown that similar settings yield an inversion efficiency of approximately 0.89 (25), which was assumed for this study. T_2_ preparation modules with hard 90º RF pulses and four adiabatic refocusing RF pulses are performed in the 8^th^, 9^th^, and 10^th^ cardiac cycles to improve T_2_ sensitization using echo times of 30, 50 and 70 ms, respectively. The Multimapping imaging parameters for all experiments are: field-of-view = 320×320 mm, spatial resolution = 2×2 mm, slice thickness = 10 mm, nominal flip angle = 50°, bandwidth = 1076 Hz/pixel, TR = 2.3 ms, TE = 1.2 ms, SENSE factor = 2, linear profile order. Ten startup RF pulses are used with linearly increasing flip angles. The Multimapping scan was acquired in a mid-ventricular short axis slice (except in one patient which was mistakenly acquired in a four-chamber view) during a breath-hold. Native Multimapping was acquired in all 47 patients, and post-contrast Multimapping was performed in 31 patients approximately 15 to 20 minutes after contrast agent administration (0.2 mmol/kg gadobutrol). Due to clinical prioritization, the post-contrast Multimapping was performed after the acquisition of post-contrast MOLLI and LGE.

**Figure 1.**
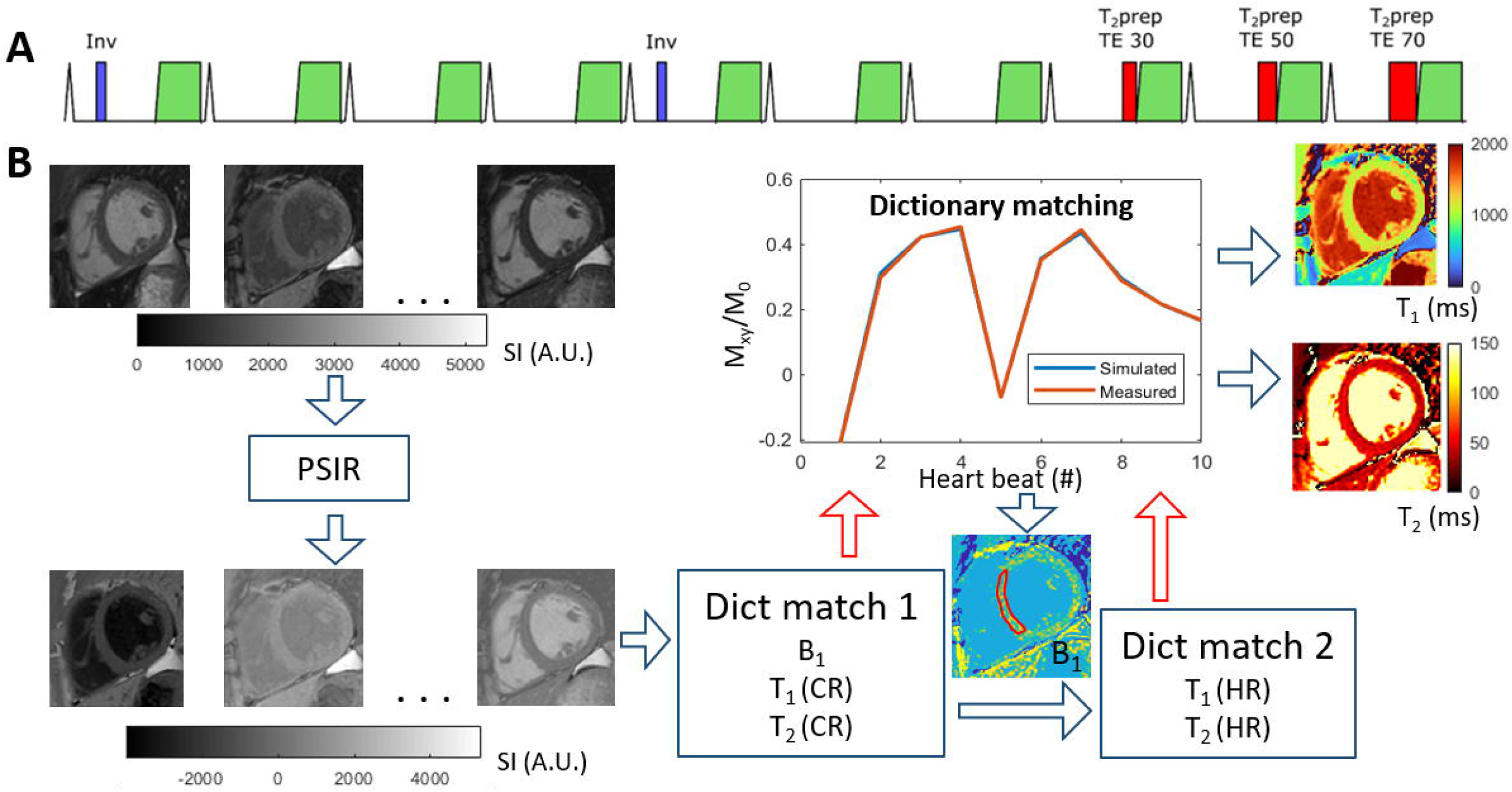
Multimapping pulse sequence (A) and post-processing (B). The pulse sequence consists of ECG-triggered 2D single-shot acquisitions in 10 consecutive cardiac cycles, preceded by inversion pulses in cycles 1 and 5, and T_2_prep with different echo times (TE) in cycles 8, 9 and 10. The post-processing involves first performing a phase sensitive inversion recovery (PSIR) operation to restore the polarity of the signal, followed by an initial dictionary matching (Dict match 1) to estimate a global B_1_ correction factor based on a ROI in the myocardial septum. The dictionary matching finds the pixel-wise closest match between acquired data and the simulated acquisition (with scan-specific delays) for different T_1_, T_2_ and RF scaling factor (B_1_) combinations. A second dictionary matching step (Dict match 2) is then performed for only T_1_ and T_2_ with high temporal granularity (1 ms resolution) to generate the final T_1_ and T_2_ maps.

All Multimapping source images were reconstructed on the scanner and transferred to an offline workstation (Intel Core i7-8565U 1.80 GHz processor with 16Gb RAM) to generate T_1_ and T_2_ maps using MATLAB R2021b (The MathWorks, Natick, MA). The MATLAB code used to generate the maps, including example Multimapping source images from one subject, is available at https://github.com/Multimapping/Matlab_files. Since blood samples were not available for all patients, Multimapping synthetic ECV maps were generated using synthetic hematocrit values, based on the native MOLLI left ventricular blood pool measurements, as previously outlined (26). Image registration using a rigid body transformation was applied to spatially align the native and post-contrast T_1_ maps prior to ECV calculation.

MOLLI was acquired in all 47 patients and T_2_bSSFP was acquired in 45 patients, in the same slice as Multimapping and used as clinical reference techniques for T_1_ and T_2_ mapping, respectively. All imaging parameters for the reference techniques (field-of-view, spatial resolution etc.) were the same as for Multimapping, except for the flip angle which was 35º. MOLLI was acquired with the 5(3s)3 scheme and used the same adiabatic inversion pulse as Multimapping. T_2_bSSFP was acquired with four images at different T_2_ preparation echo times (0, 23, 46 and 70 ms) and used 3 pause cardiac cycles between each image. Furthermore, T_2_bSSFP used the same RF pulse types for the T_2_ preparation module as Multimapping. The reference maps were reconstructed on the scanner using vendor-provided inline mapping algorithms, except for the ECV maps which were generated offline using MATLAB. Similar to Multimapping, synthetic ECV maps were generated using a synthetic hematocrit value derived from the left ventricular blood pool T_1_ measured in the native MOLLI image. As for Multimapping, image registration was applied to native and post-contrast MOLLI T_1_ maps before ECV was calculated. LGE imaging parameters were TR/TE = 5.6/2.0 ms, flip angle = 25º, FOV = 350×350 mm^2^, spatial resolution = 1.8×1.8 mm^2^.

### 2.3 Image Analysis

T_1_ (native and post-contrast), T_2_ (native) and ECV measurements were made by drawing manual regions-of-interest (ROI) in all datasets. To compare Multimapping to the clinical reference techniques, two sets of myocardial measurements were performed, one targeting any diseased myocardium and one targeting healthy myocardium. For the measurements of diseased myocardium in all maps, ROIs were drawn in the area corresponding with the most prominent positive LGE findings of each patient. Since only a subset of patients had positive LGE findings in the imaged slice, this resulted in 21 measurements for native T_1_ and T_2_, and 12 measurements for post-contrast T_1_ and ECV. For the measurements of healthy myocardium in all maps, ROIs were drawn in the area remote of any LGE abnormality and preferentially in the interventricular septum if it was free of abnormal LGE. Patients were excluded from this analysis if there were indications suggestive of global or diffuse myocardial disease. The measurements in the healthy myocardium were performed in a total of 19 patients for native T_1_, 12 patients for T_1_ post-contrast and ECV, and 18 patients for T_2_.

Measurements were performed in all patients by one observer (CJ, 1 year of CMR experience). To allow intra-observer variability analysis, measurements were repeated in 23 patients by the same observer two weeks later. For inter-observer variability analysis, the same 23 patients were measured by two additional observers (MH and CJC with 14 and 21 years of CMR experience, respectively). Furthermore, to compare blood T_1_ (native and post-contrast), ROIs were drawn in the left ventricular blood pool (avoiding any papillary muscles) in the Multimapping T_1_ and MOLLI images.

The image quality of the acquired maps were qualitatively compared using a Likert scale as devised by Jaubert et al. (22) with the following categories: 1 = uninterpretable, 2 = poor definition of edges, significant noise and/or residual artifacts, 3 = mildly blurred edges, mild noise and/or residual artifacts, 4 = slightly blurred edges, minor residual artifacts, 5 = negligible blurring or residual artifacts. Visual scoring was performed for T_1_ (native and post-contrast), T_2_ (native) and ECV separately using the different mapping techniques and this analysis was performed in 20 patients. The visual scoring was performed by consensus of two blinded observers (CJ and CJC).

### 2.4 Statistical Analysis

Continuous variables are expressed as mean ± SD. Categorical variables are expressed as counts and percentages. For the remote measurements, two-tailed Student paired t-tests were performed to compare Multimapping to MOLLI for native T_1_, post-contrast T_1_ and ECV, and Multimapping and T_2_bSSFP for native T_2_. For the remote measurements, all parameters tested positive for normality using a Shapiro-Wilk test. Bland-Altman and correlation plots were used to evaluate the agreement and correlation, respectively, between Multimapping and the reference techniques of the measurements in diseased myocardium for native T_1_, post-contrast T_1_, ECV and native T_2_. To investigate any heart-rate dependency for the mapping techniques, the measurements of the remote myocardium was correlated with the heart-rate at the time of the scan. Similarly, dependency on T_2_ for T_1_ (and vice-versa) was evaluated by correlating remote T_1_ with T_2_ for both Multimapping and the reference techniques, and testing for statistical significance. To account for multiple comparisons, Bonferroni correction was performed on the threshold for all significance tests. Since four comparisons were performed (native and post-contrast T_1_, native T_2_ and ECV), a threshold of 0.05/4 = 0.0125 was used. Intra-observer repeatability and inter-observer repeatability was assessed with intraclass correlation coefficient (ICC) analysis. ICC was calculated using absolute agreement two-way mixed model. Statistical analysis was performed using IBM SPSS Statistics, version 27.0.

## 3 Results

Representative parameter maps acquired with Multimapping, reference techniques and LGE in a patient with no cardiovascular disease findings are shown in Figure 2. Parameter maps from a patient with myocarditis are shown in Figure 3, with prominently altered quantitative values seen in both Multimapping and reference techniques. The final example, in Figure 4, shows parameter maps from a patient with myocardial infarction with a clearly delineated area of infarction in the Multimaps, correlating with LGE. Multimaps for all patients can be downloaded from https://github.com/Multimapping/Patient_study/raw/main/MapReconstructions.pdf.

**Figure 2.**
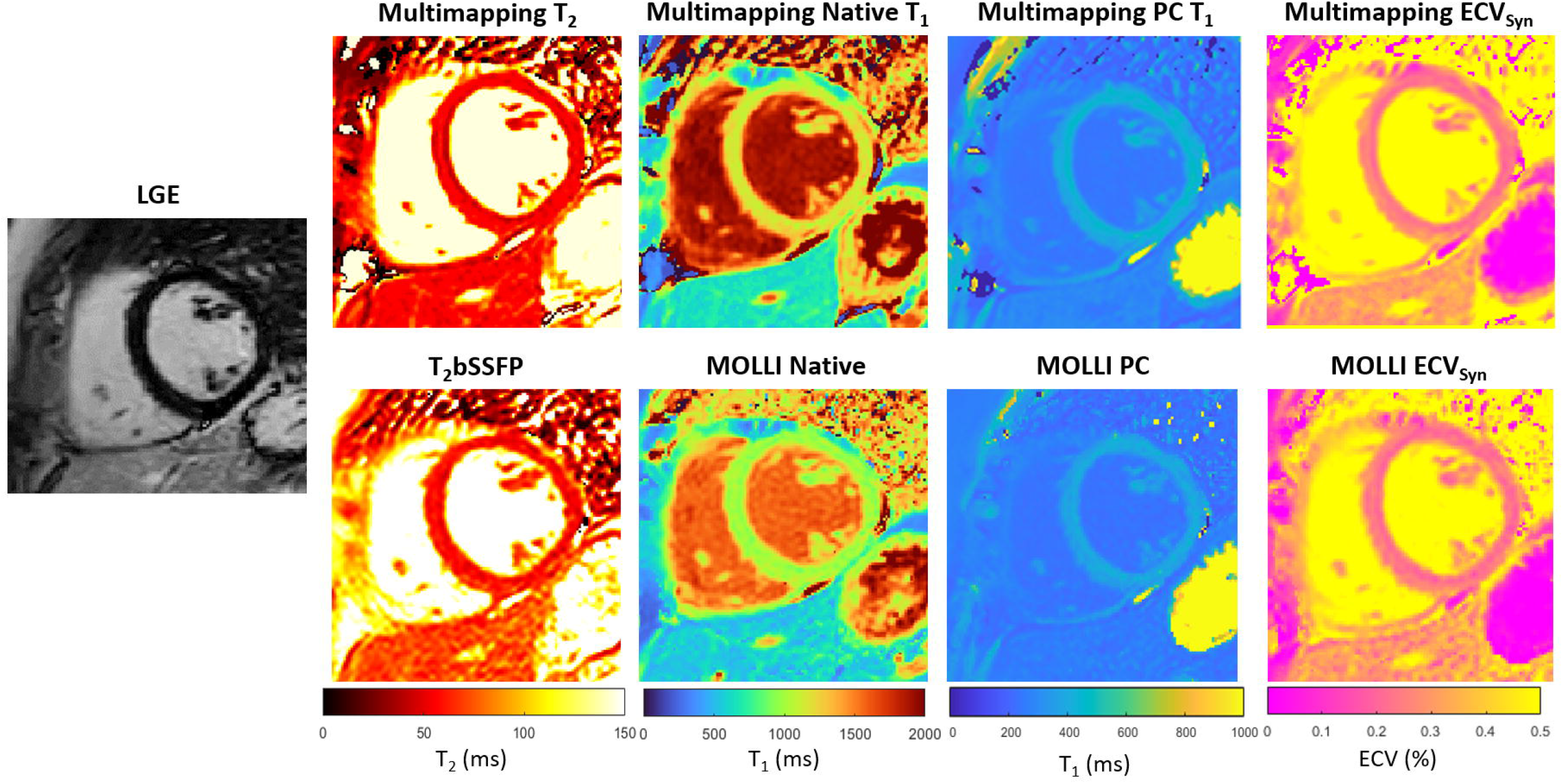
LGE and parameter maps (native T_2_, native T_1_, post-contrast T_1_ and ECV) are shown for a patient with no cardiovascular disease findings. The parameter maps were acquired using either Multimapping (top row) or reference techniques (bottom row). Septal T_2_ was 49.6 ms and 56.7 ms for Multimapping and T_2_bSSFP, respectively. Septal (native/post-contrast) T_1_ was 1144/422 ms and 1003/381 ms for Multimapping and MOLLI, respectively. Septal ECV was 24.9% and 23.8% for Multimapping and MOLLI, respectively.

**Figure 3.**
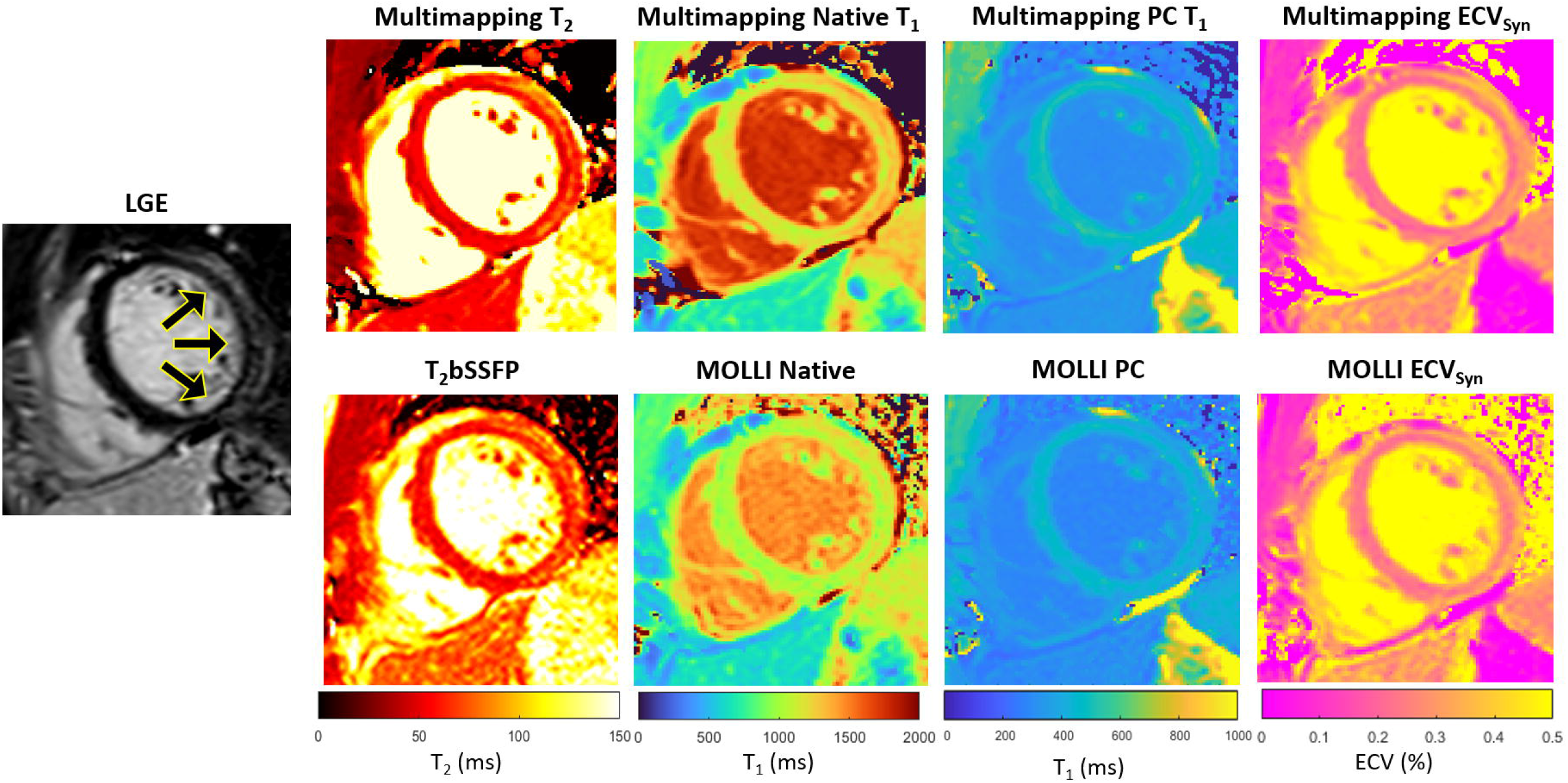
LGE and parameter maps in a patient with myocarditis, as indicated by the increased signal in the lateral wall in the LGE and also apparent as altered values in the parameter maps (Multimapping top row, reference techniques bottom row). Measurements in the area of enhancement (lateral wall) yielded T_2_ of 65.7 ms and 62.4.6 ms for Multimapping and T_2_bSSFP, respectively. T_1_ values (native/post-contrast) in the same area were 1286/446 ms and 1111/423 ms for Multimapping and MOLLI, respectively. ECV was 26.6% and 28.3% in the enhanced area for Multimapping and MOLLI, respectively.

**Figure 4.**
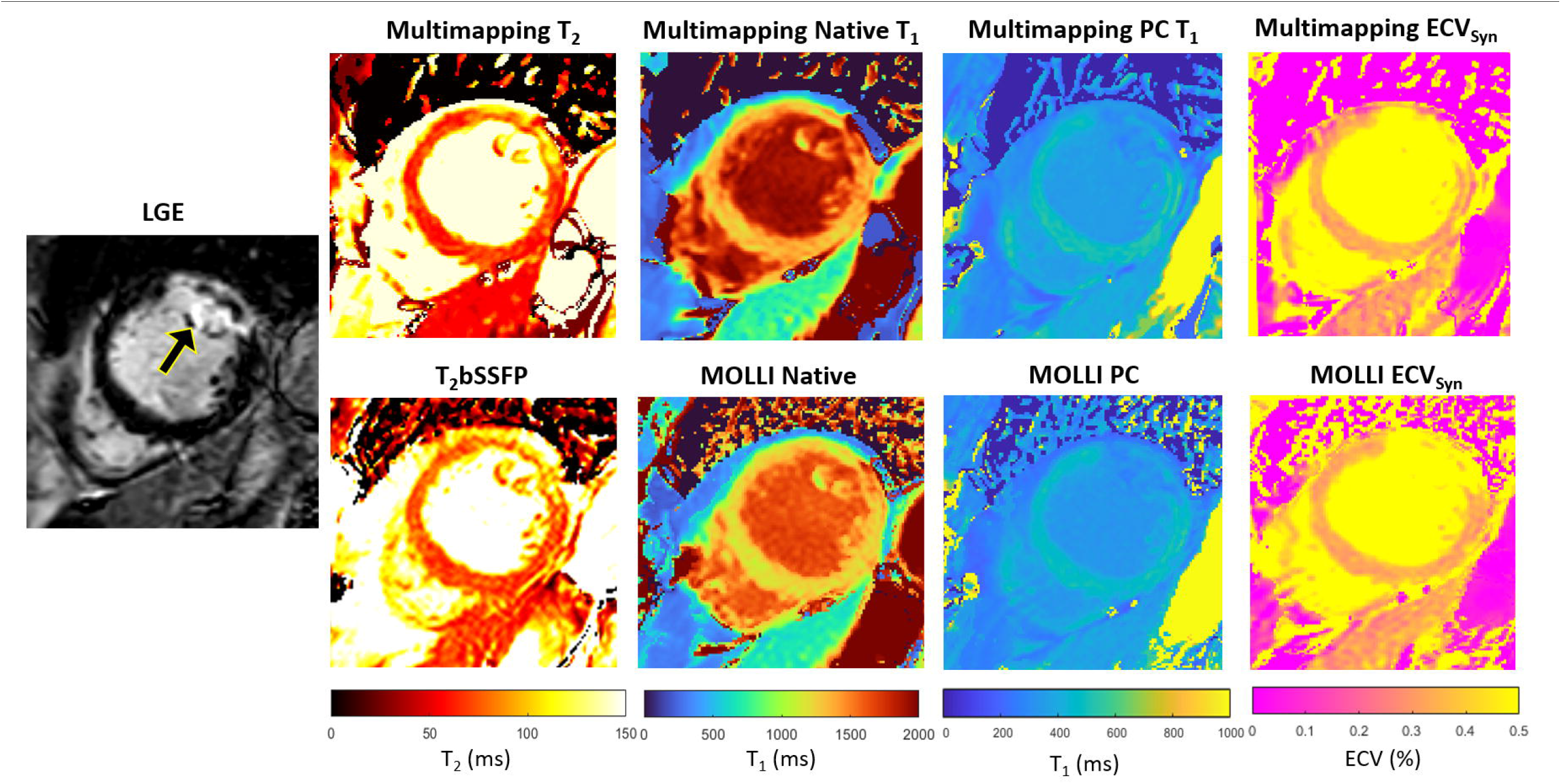
LGE and parameter maps in a patient with myocardial infarction, clearly visualized using all techniques, both native and post-contrast. Measurements in the area of enhancement (anterolateral segment) yielded T_2_ of 90.5 ms and 88.6 ms for Multimapping and T_2_bSSFP, respectively. T_1_ values (native/post-contrast) in the same area were 1516/355 ms and 1440/338 ms for Multimapping and MOLLI, respectively. ECV was 33.3% and 33% in the enhanced area for Multimapping and MOLLI, respectively.

### 3.1 Comparison of Multimapping and reference techniques in remote myocardium

The Multimapping native T_1_ was 1116±21 ms and for MOLLI 1002±21 ms, resulting in a statistically significant bias of 114 ms (*P* < 0.001). Multimapping post-contrast T1 was 479±31 ms and for MOLLI 426±27 ms, yielding a bias of 53 ms which was statistically significant (*P* < 0.001). Multimapping ECV was 21.5±1.9%, MOLLI ECV was 23.7±2.3%, resulting in a bias of -2.2% which was statistically significant (*P* = 0.001). Multimapping native T_2_ was 48.0±3.0 ms while T_2_bSSFP was 53.9±3.5 ms, a statistically significant bias of -5.9 ms (*P* < 0.001) (Figure 5).

**Figure 5.**
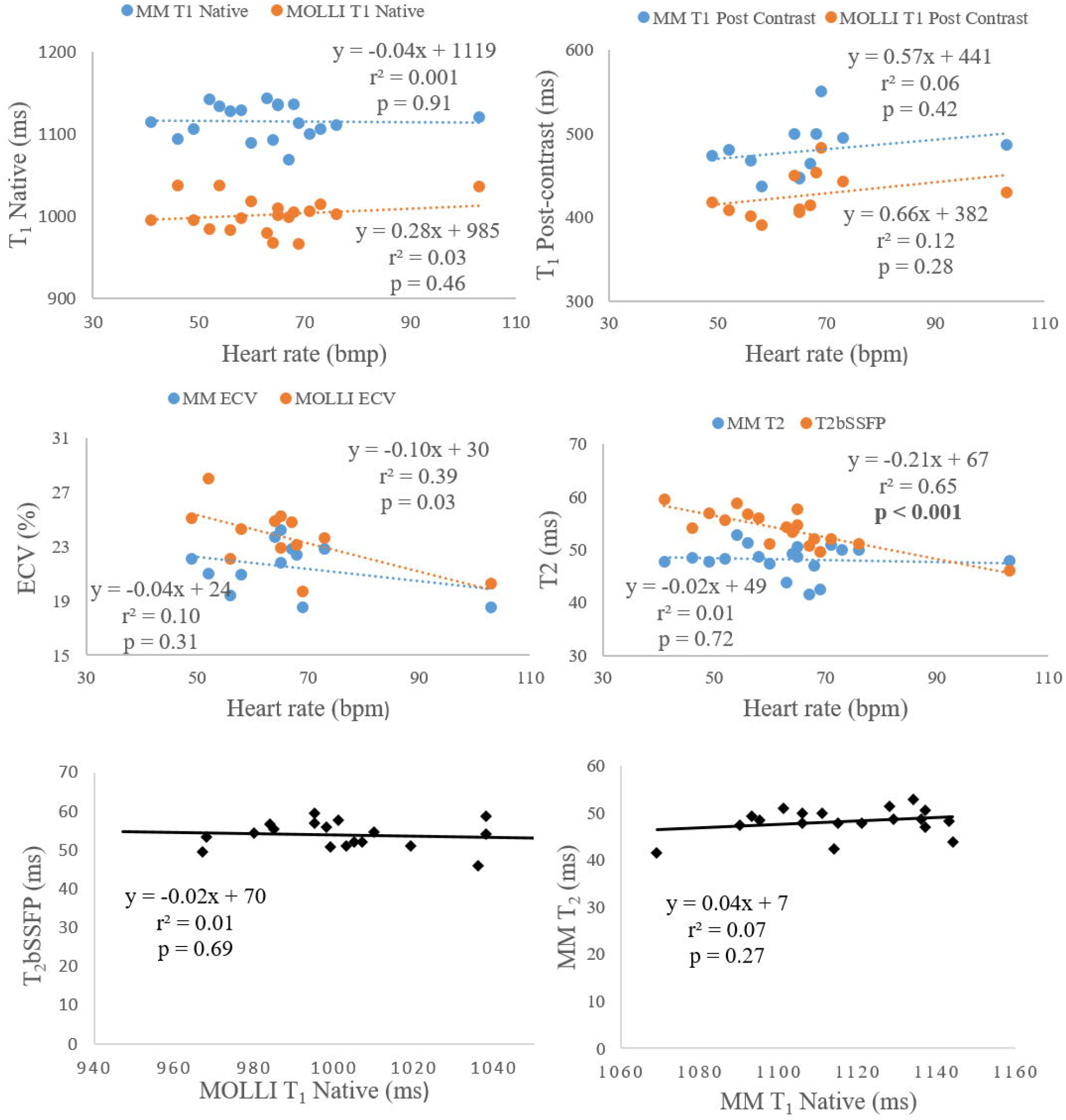
The two top rows show the correlation analysis of heart-rate dependency to T_1_ (native and post-contrast), ECV and T_2_. The bottom row shows the correlation analysis between native T_1_ and T_2_.

There was no correlation between native T_1_ and T_2_ for neither Multimapping nor MOLLI and T_2_bSSFP. Multimapping T_1_ (native and post-contrast), T_2_ or ECV and MOLLI T_1_ (native and post-contrast) or ECV did not correlate with heart-rate either. However, T_2_bSSFP showed a correlation with heart rate (*P* < 0.001) (Figure 5).

### 3.2 Comparison of Multimapping and reference techniques for diseased myocardium

In general, the correlation between Multimapping and the clinical reference techniques was very strong (r^2^ > 0.7) for most variables (Figure 6). A strong correlation coefficient (r^2^ > 0.5) was found between Multimapping and MOLLI for myocardial T_1_ post-contrast (r^2^ = 0.66) and blood T_1_ post-contrast (r^2^ = 0.53).

**Figure 6.**
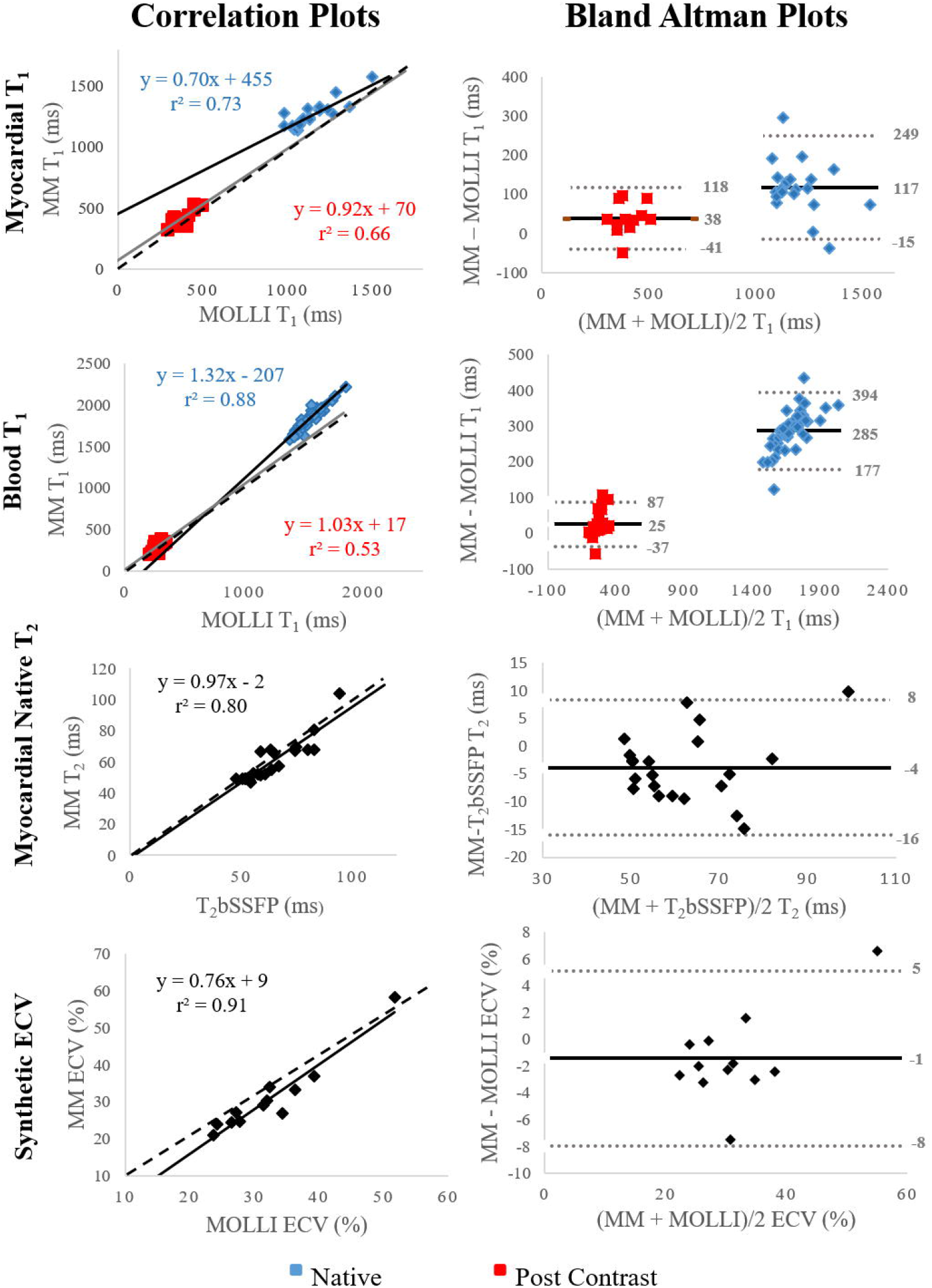
Correlation and Bland-Altman plots comparing Multimapping (MM) to MOLLI and T_2_bSSFP. For the correlation plots, black and grey lines indicate line of best fit, and the dottet lines show the identity lines. The black lines indicate bias in the Bland-Altman plots. Correlation coefficient (r^2^) is reported in the correlation plots and mean difference, lower and upper limits of agreement (1.96 SD) in the Bland-Altman plots.

### 3.3 Inter- and intra-observer variability

The myocardial measurements and measurements of the left ventricular blood pool for intra- repeatability assessment showed excellent repeatability (myocardial ICC > 0.97, LV blood pool ICC = 1.00) (Table 2). The myocardial measurements for inter-repeatability showed moderate to excellent repeatability (ICC > 0.73) for all mapping techniques. The native and post-contrast T_1_ measurements of the blood pool for inter-repeatability showed good to excellent repeatability (ICC > 0.92).

**Table 2.** Inter and intra-observer ICC (95% confidence interval).

|  | Myocardium |  | Blood pool |  |
| --- | --- | --- | --- | --- |
|  | Intra-repeatability | Inter-repeatability | Intra-repeatability | Inter-repeatability |
| MM T<sub>1</sub> Native | 0.99 (0.97-1.00) | 0.87 (0.76-0.94) | 1.00 (1.00-1.00) | 0.92 (0.85-0.96) |
| MOLLI T<sub>1</sub> Native | 0.99 (0.97-0.99) | 0.93 (0.88-0.97) | 1.00 (1.00-1.00) | 0.97 (1.00-1.00) |
| MM T<sub>1</sub> PC | 0.99 (0.98-1.00) | 0.73 (0.54-0.86) | 1.00 (1.00-1.00) | 1.00 (1.00-1.00) |
| MOLLI T<sub>1</sub> PC | 0.97 (0.93-0.99) | 0.95 (0.90-0.98) | 1.00 (1.00-1.00) | 1.00 (1.00-1.00) |
| MM ECV | 0.99 (0.98-1.00) | 0.94 (0.89-0.97) |  |  |
| MOLLI ECV | 0.99 (0.97-0.99) | 0.93 (0.87-0.97) |  |  |
| MM T<sub>2</sub> | 0.99 (0.96-0.99) | 0.91 (0.82-0.95) |  |  |
| T<sub>2</sub>bSSFP | 0.98 (0.94-0.99) | 0.88 (0.78-0.95) |  |  |

MM, Multimapping; ECV, extracellular volume fraction; MOLLI, modified Look-Locker inversion recovery; T<sub>2</sub>bSSFP, T<sub>2</sub> prepared balanced steady-state free precession.

### 3.4 Image quality assessment

The image quality was scored significantly higher for Multimapping compared to T_2_bSSFP (*P* < 0.001), MOLLI native T_1_ (*P* = 0.007), MOLLI post-contrast T_1_ (*P* < 0.001), and MOLLI ECV (*P* < 0.001) (Figure 7).

**Figure 7.**
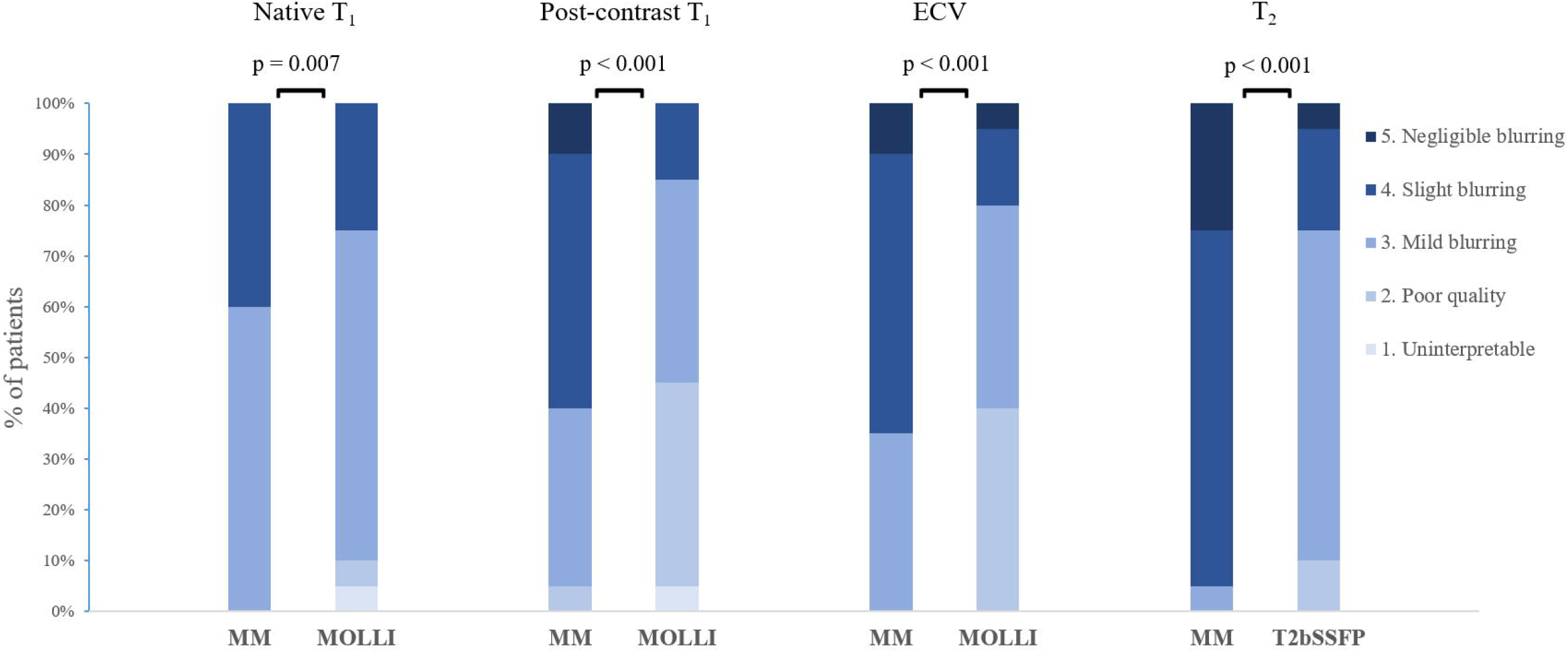
Distribution of image quality scores for Multimapping (MM) and in-vivo reference techniques.

## 4 Discussion

In this study, a new method for simultaneous T_1_ and T_2_ mapping was compared to clinical reference mapping technique in a cohort of patients with cardiovascular disease. We found a strong to very strong correlation between the methods for all measured parameters (native T_1_, post-contrast T_1_, ECV and native T_2_), while the image quality was considered better using the proposed Multimapping technique compared to the reference methods. Furthermore, intra- and inter-observer variability of Multimapping parameter measurements were in general low and similar to those obtained with the clinical reference techniques.

In segments of healthy/remote myocardium we measured a mean native T_1_ of 1116 ms using Multimapping, more than 100 ms higher than for MOLLI. However, MOLLI is known to significantly underestimate T_1_ when compared to more accurate methods such as SASHA (5), which typically yields native T_1_ of around 1200 ms at 1.5T (7,27). The native T_1_ Multimapping values are also in line with the previous study using this technique in healthy volunteers which measured 1114 ms (24). For post-contrast T_1_ mapping there was also a significantly longer T_1_ using Multimapping (479 ms) compared to MOLLI (427 ms). Although post-contrast T_1_ values are more difficult to compare between studies due to differences in contrast agents and the timing of acquisition after injection, previous studies have shown underestimation of post-contrast T_1_ for MOLLI compared to more accurate techniques such as SASHA (28,29). The study by Nordlund et al. (29) also demonstrated that MOLLI overestimates ECV by approximately 4% in healthy volunteers compared to SASHA, the latter technique correlating more closely with radioisotopes in pigs. This suggests that the significantly lower ECV measured in this study with Multimapping (22%) may be more accurate compared to MOLLI (24%). However, the conversion from T_1_ to hematocrit was based on the relationship established for MOLLI in a previous study, which may bias measurements if applied to Multimapping synthetic ECV. For Multimapping synthetic ECV to be used independently of MOLLI then the relationship between Multimapping blood T_1_ and hematocrit should be established. Alternatively, the hematocrit could be directly measured to calculate Multimapping ECV without the need for a MOLLI acquisition. Correlation of T_1_ and T_2_ values with potential confounding variables such as heart-rate or the opposite (T_2_ or T_1_) parameter did not show any particular dependency for Multimapping in this regard. However, T_2_bSSFP appeared to be inversely correlated with heart-rate. This suggest additional delay cardiac cycles may be required to yield less biased T_2_ values for high heart-rates. Conversely, Multimapping may be a more robust approach for T_1_ and T_2_ mapping at higher heart rates as no additional modification of the pulse sequence is required.

In the measurements of myocardial segments with disease, we found a high correlation between Multimapping and the clinical reference techniques for native T_1_ (blood and myocardium), T_2_, and ECV. While correlations for post-contrast T_1_ (blood and myocardium) was more moderate, this may be explained by the confounding factor of time after injection, which affect the T_1_ measurements. Furthermore, measured post-contrast T_1_ in this study had a narrower range for both Multimapping and MOLLI compared to native T_1_ which can contribute to a weaker correlation between techniques. Nevertheless, a very strong correlation between Multimapping and reference techniques for native T_1_, T_2_ and ECV indicates that Multimapping is a useful technique that can be used to detect disease.

Dictionary-based mapping techniques such as Multimapping typically assumes that there is no through-plane motion, which is not the case for flowing blood. Such through-plane motion leads to T_1_ over-estimation as inflowing spins has seen fewer RF pulses and is therefore less saturated. This can explain the observed over-estimation of blood (particularly native) T_1_ relative to MOLLI. However, it should also be noted that, due to the strong correlation for native blood T_1_ blood between Multimapping and MOLLI, the Multimapping technique can likely still capture variability in blood T_1_ (due to e.g. different hematocrit levels) with a similar sensitivity as MOLLI.

The image quality was superior using Multimapping compared to all clinical reference techniques. This could be due to the higher flip angle of 50º using Multimapping, compared to MOLLI and T_2_bSSFP which both use a flip angle 35º, with otherwise identical imaging parameters to Multimapping. A higher flip angle for bSSFP-based mapping techniques leads to a higher signal-to-noise ratio which typically contributes to improved image quality. The shorter duration of Multimapping (10 beats) compared to both MOLLI (11 beats) and T_2_bSSFP (16 beats) means that Multimapping is less prone to respiratory motion-induced misalignment, which may also contribute to a better image quality. While the Multimapping and MOLLI pulse sequences are very similar (both inversion recovery with bSSFP readouts), Multimapping benefits from phase sensitive inversion recovery post-processing step which has been shown to improve T_1_ map image quality compared to fitting with magnitude images (30), used in the vendor provided fitting algorithm for MOLLI. Compared to Multimapping, T_2_bSSFP uses significantly fewer source images for T_2_ mapping, and while only three T_2_ preparation modules are included in the Multimapping pulse sequence, the bSSFP readout is intrinsically T_2_/T_1_ weighted which contributes to the T_2_ sensitivity and may explain the improved image quality.

The intra- and inter-observer repeatability analysis showed an excellent repeatability for most measurements using both Multimapping and reference techniques. While Multimapping post-contrast myocardial T_1_ inter-observer ICC of 0.73 was relatively low compared to that of MOLLI (ICC = 0.95), post-contrast T_1_ mapping is primarily used to generate ECV maps, and here Multimapping and MOLLI yielded near identical ICC values of 0.94 and 0.93, respectively.

### 4.1 Comparison with other simultaneous T_1_ and T_2_ mapping techniques

Several simultaneous T_1_ and T_2_ mapping techniques have been proposed over the last years, comparable to Multimapping. Published studies using similar methods in healthy volunteers are summarized in Table 3, including Multimapping (24). Multimapping has a shorter scan duration than nearly all other simultaneous T_1_ and T_2_ mapping techniques, requiring 10 beats, which is also shorter than both MOLLI and T_2_bSSFP. As many patients with cardiovascular diseases struggle to hold their breath for an extended period, reducing the scan time of mapping techniques is important and has been the focus of several studies (11,31). This is also in line with the endeavor of utilizing less time-consuming CMR protocols in order to improve adoption of CMR in routine cardiovascular practice. Inversion recovery magnetization preparation pulses are often used for myocardial T_1_ mapping as they increase quantification precision compared to saturation recovery (5), using the full dynamic range of the longitudinal magnetization, at the expense of accuracy as inversion pulses are more sensitive to confounding elements such as magnetization transfer and transverse relaxation during the pulses, which reduce their efficiency (6,25,32). Therefore, saturation recovery technique measurements are generally considered to be closer to the “true” in-vivo T_1_ times, typically several 100 ms higher than MOLLI on either 1.5T and 3T scanners. In this regard, Multimapping which uses inversion recovery, generates T_1_ values in healthy/remote myocardium of 1116 ms, which is closer to the saturation recovery based techniques (of approximately 1200 ms) than MOLLI (approximately 1000 ms) or the most comparable simultaneous T_1_ and T_2_ mapping studies, Blume et al. (15) and Jaubert et al. (33), which report a myocardial T_1_ of 1017 ms and 1045 ms, respectively. This may be due to the assumed lower inversion efficiency of 0.89 for the inversion pulses, incorporated into the Multimapping signal model, which is likely closer to the true inversion efficiency than assuming perfect efficiency. However, the inversion efficiency potentially varies between field strengths and vendors, or even spatially across an image due to B_0_ and B_1_ inhomogeneity. Furthermore, the current Multimapping technique does not consider magnetization transfer. To yield more accurate T_1_ values, reproducible across scanner platforms, these confounding effects should be included in the Multimapping signal model, preferably on a pixel-wise basis, although this will likely negatively impact the precision.

**Table 3.** Comparable published simultaneous T_1_ and T_2_ mapping techniques.

| Study | Scan time | FB/BH | IR/SR | Readout | Subjects | Additional mapping | Field strength | Native T1 (ms) | Native T2 (ms) |
| --- | --- | --- | --- | --- | --- | --- | --- | --- | --- |
| Blume et al. (15) | Around 3 minutes | FB | IR | 2D Cartesian bSSFP | 19 HV | - | 1.5T | 1017±91 | 50±4 |
| Kvernby et al. (17) | 15 beats | BH | IR | 3D Cartesian GRE | 10 HV | - | 3T | 1083±43 | 50.4±3.6 |
| Akçakaya et al. (18) | 13 beats | BH | SR | 2D Cartesian bSSFP | 10 HV | - | 1.5T | 1210±24 | 48.2±2.8 |
| Santini et al. (35) | 8 beats | BH | IR | 2D Cartesian bSSFP | 5 HV | - | 3T | 1227±68 | 37.9±2.4 |
| Hamilton et al. (36) | 15 beats | BH | IR | 2D spiral GRE | 11 HV | - | 3T | 1235, range 1199 - 1316 | 38, range 32 – 43 |
| Jaubert et al. (33) | 15 beats | BH | IR | 2D radial ME-GRE | 10 HV | PDFF | 1.5T | 1045±32 | 42.8±2.6 |
| Christodoulou et al. (19) | 88 seconds | FB | IR | 2D radial GRE | 10 HV | Cardiac motion | 3T | 1216±67 | 47.8±4.9 |
| Shao et al. (37) | 11 beats | BH | IR | 2D radial GRE | 10 HV | - | 3T | 1366±31 | 37.4±0.9 |
| Guo et al. (38) | 1.4±0.3 minutes (WH) | FB | SR | M2D Cartesian bSSFP | 13 HV | - | 3T | 1373±50 | 48.7±2.5 |
| Hermann et al. (39) | 18.5 seconds (+resp gating) | FB | SR | 2D Cartesian ME-GRE | 10 HV | T2* | 3T | 1573±86 | 33.2±3.6 |
| Chow et al. (40) | 11 beats | BH | SR | 2D Cartesian bSSFP | 10 HV | - | 3T | 1523±18 | 36.7±1.1 |
| Jarkman et al. | 10 beats | BH | IR | 2D Cartesian bSSFP | Remote myocardium 19 patients | - | 1.5T | 1116±21 | 48.0±3.0 |

It can be difficult to precisely pinpoint the sources of differences in T_1_ and T_2_ values for the different techniques outlined in Table 3, particularly as many techniques rely on unconventional acquisition, reconstruction and mapping strategies. These include, for example, non-Cartesian (radial or spiral) trajectories with iterative reconstruction algorithm, coupled with sophisticated and advanced mapping techniques which may be difficult to reproduce. In contrast, the Multimapping pulse sequence consists of a MOLLI-like acquisition scheme (inversion recovery with Cartesian single-shot 2D bSSFP readout) which are available on all major vendor platforms, with the addition of T_2_prep modules which have also been implemented on all vendor platforms. For transparency, the Multimapping parameter mapping method using dictionary matching has been provided open source to enable reproduction of this technique by other investigators which may also enable direct comparison of Multimapping with other simultaneous T_1_ and T_2_ mapping techniques.

### 4.2 Limitations

This study has several limitations: no respiratory motion correction was performed. Correcting for respiratory-induced image misalignment is important even for breath-held scans and can be achieved using image registration (34). Although image registration could be readily applied to Multimapping source image to this end, this was not performed in order to have a fair comparison with MOLLI and T_2_bSSFP maps which were generated using inline vendor algorithm without motion correction. A second technical limitation of Multimapping is that manual interaction is required to define a ROI in the myocardial septum for the B_1_ estimation. However, this is a relatively simple step, comparable to the input required to define ROIs for ECV maps. Further work is required to automatize this step or to incorporate B_1_ in the high resolution T_1_ and T_2_ dictionary matching, which would obviate the need for any manual interaction but with a potential penalty to the precision. A study limitation is that the patient cohort consisted of a small, heterogeneous population of patients with various cardiovascular diseases, and performed on a single 1.5T Philips scanner. Further studies are required to evaluate the performance of Multimapping at 3T and using other vendors. The evaluation of heart-rate dependence of the mapping techniques was limited by the relatively narrow heart rates of nearly all patients (only one with heart-rate over 100 bpm).

## 5 Conclusions

Multimapping T_1_ and T_2_ values show high correlations with clinical reference mapping techniques in a diverse cohort of patients with different cardiovascular diseases. Multimapping enables simultaneous T_1_ and T_2_ mapping and can be performed in a short (10 cardiac beats) breath-hold, with image quality superior to that of the clinical reference techniques.

## 7 Author Contributions

MH and CJC conceived of the study. MH developed the methods, acquired the data, performed image reconstruction and processing. MH, CJ, and CJC analyzed the data. All authors participated in revising the manuscript, read and approved the final manuscript.

## 9 Acknowledgments

Not applicable.

## 10 Data Availability Statement

The Multimapping post-processing pipeline (Matlab code) can be downloaded from: https://github.com/Multimapping/Matlab_files. Images for all patients included in this study can be downloaded from: https://github.com/Multimapping/Patient_study/raw/main/MapReconstructions.pdf.

